# Four-sphere head model for EEG signals revisited

**DOI:** 10.1101/124875

**Authors:** Solveig Næss, Chaitanya Chintaluri, Torbjørn V. Ness, Anders M. Dale, Gaute T. Einevoll, Daniel K. Wójcik

**Author notes:** Equal contribution (Solveig Næss, Chaitanya Chintaluri).

## Abstract

Electric potential recorded at the scalp (EEG) is dominated by contributions from current dipoles set by active neurons in the cortex. Estimation of these currents, called ’inverse modeling’, requires a ’forward’ model, which gives the potential when the positions, sizes, and directions of the current dipoles are known. Different models of varying complexity and realism are used in the field. An important analytical example is the *four-sphere model* which assumes a four-layered spherical head where the layers represent brain tissue, cerebrospinal fluid (CSF), skull, and scalp, respectively. This model has been used extensively in the analysis of EEG recordings. Since it is analytical, it can also serve as a benchmark against which numerical schemes, such as the Finite Element Method (FEM), can be tested. While conceptually clear, the mathematical expression for the scalp potentials in the four-sphere model is quite cumbersome, and we observed the formulas presented in the literature to contain errors. We here derive and present the correct analytical formulas for future reference. They are compared with the results of FEM simulations of four-sphere model. We also provide scripts for computing EEG potentials in this model with the correct analytical formula and using FEM.

## 1 Introduction

Electroencephalography (EEG), that is, the recording of electrical potentials at the scalp, has been of key importance for probing human brain activity for more than half a century (Schomer and da Silva, 2012). It is common to interpret the EEG signal in terms of current dipoles set up by active neurons (Hämäläinen et al., 1993; Sanei and Chambers, 2007). Estimation of the underlying sources based on EEG signals is called *inverse modeling*, and its key ingredient is a *forward model* for computation of the resulting signal from known current sources. While the link between the current sources and the resulting potentials in principle is well described by volume-conductor theory, the practical application of this theory is not easy because the cortical tissue, the cerebrospinal fluid (CSF), the skull, and the scalp, all have different electrical conductivities (Nunez and Srinivasan, 2006).

An important analytical forward model is the *four-sphere model* (Srinivasan et al., 1998; Nunez and Srinivasan, 2006) assuming a four-layered spherical head model where the four layers represent brain tissue, CSF, skull, and scalp, respectively. The Poisson equation, which describes the electric fields of the brain within volume-conductor theory, is solved for each layer separately, and the mathematical solutions are matched at the layer interfaces to obtain an analytical expression for the EEG signal as set up by a current dipole in the brain tissue. This model has been extensively used in analysis of EEG signals, see, e.g., Peraza et al. (2012); Wong et al. (2008); Chu et al. (2012), but it is also useful for validation of general numerical schemes, such as the Finite Element Method (FEM) (Larson and Bengzon, 2013). The FEM approach is not limited by specific assumptions on head symmetry and can, in principle, take into account an arbitrarily complex spatial distribution of electrical conductivity representing the electrical properties of the head (Bangera et al., 2010; Huang et al., 2016). This is done by building a numerical mesh for the head model with the electrical conductivity specified at each mesh point. The mesh construction is a research problem by itself and several mesh-generation tools are available, which often provide slightly different results (Geuzaine, 2009; Kehlet, 2016). The analytical solution for the four-sphere model can thus serve as a ground-truth against which an FEM implementation can be validated.

While conceptually clear, the mathematical expression of the four-sphere forward model is quite involved, and rederiving the expression we discovered errors in the formulas both in the original paper, Srinivasan et al. (1998), and in the classic EEG reference book, Nunez and Srinivasan (2006). As a consequence, the listed formulas predict incorrect EEG scalp potentials. Due to the importance of the four-sphere model, here we derive and present the correct analytical formulas for future reference. We further show that this formula, unlike the previous ones, gives predictions in accordance with FEM simulations.

To facilitate its use in further research we also provide numerical scripts for computing EEG potentials with the corrected formulas, as well as FEM simulation code.

## 2 Methods

### 2.1 Four-sphere model

The well-established volume-conductor theory is based on the quasi-static approximation to the Maxwell’s equations. The electric potential Φ is found here by solving Poisson’s equation (Nunez and Srinivasan, 2006),

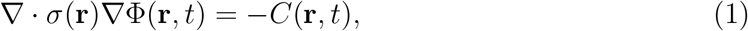

where *C*(**r**, *t*) is the density of current sources. *σ*(**r**) is the position-dependent conductivity of the medium, here assumed to be isotropic so that *σ*(**r**) is a scalar. The four-sphere model is a specific solution of this equation which assumes that the conductive medium consists of four spherical layers representing specific constituents of the head: brain tissue, CSF, skull, and scalp (Figure 1A). In the computations below, these layers are labeled by *s* = 1 to 4, respectively. The conductivity *σ_s_*(**r**) is assumed to be homogeneous, i.e., constant within each layer and independent of frequency (Pettersen et al., 2012). In the examples below we assume the same values of conductivities and concentric shell radii as in Nunez and Srinivasan (2006), see Table 1. The solution of Equation (1) is subject to the following boundary conditions (where *s* = 1, 2, 3), assuring continuity of both electrical potential and current across the layer boundaries, and no current escaping the outer layer (Nunez and Srinivasan, 2006):

**Figure 1:**
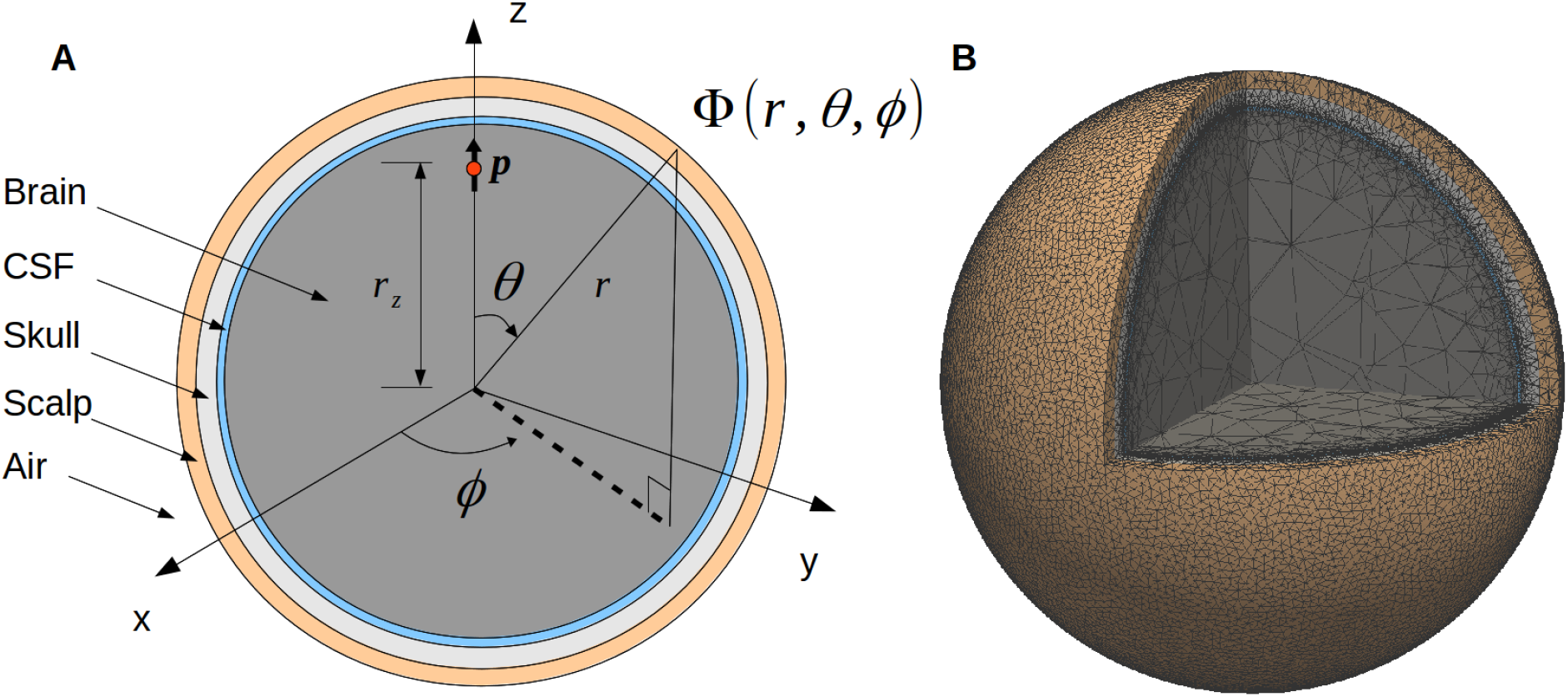
Illustration of the four-sphere head model. **(A)** Cross-section of the four-sphere head model, with the different colors corresponding to the different head layers: brain, CSF, skull, and scalp. The current dipole **p** is located in the brain layer, at a distance *r_z_* from the center of the sphere. In all the subsequent figures, the dipole is placed in the *x* = 0 plane, at the z-axis (*r_z_* = 7.8 cm). **(B)** Mesh of the four-sphere model used in the FEM simulations illustrating the different electrical conductivity values for each of the spheres.

**Table 1:** Radii and electrical conductivities of the present four-sphere model. *σ* is the conductivity in each of the specified regions. Three variants of the model were considered with skull conductivity reduced by a factor *K* (20, 40, or 80) compared to the conductivity of the brain.

| Labels | Name | Radius (cm) | $\sigma$ (S/m) |
| --- | --- | --- | --- |
| 1 | Brain | 7.9 | 0.33 |
| 2 | CSF | 8.0 | $5 \sigma_{\text{brain}}$ |
| 3 | Skull | 8.5 | $\sigma_{\text{brain}}/K$ |
| 4 | Scalp | 9.0 | $\sigma_{\text{brain}}$ |

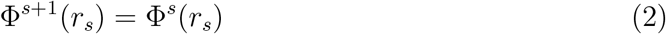

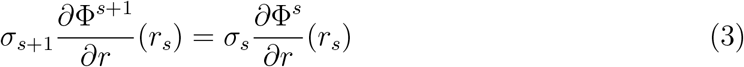

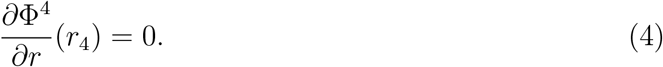

### 2.2 Analytical solution of the four-sphere head model

The solution of Equation (1) takes different forms for tangential and radial dipoles, and any dipole can be decomposed into a linear combination of these two. The following derivations are based on Appendix G and H in Nunez and Srinivasan (2006), and are described in more detail in Appendix A.

#### 2.2.1 Radial dipole

Nunez and Srinivasan (2006) give the following equations for calculating extracellular potentials from a radial dipole in the four-sphere model: The potential in the inner sphere, the brain, is given by Φ^1^(*r, θ*), while Φ^*s*^(*r, θ*) gives the potential in CSF, skull, and scalp, for *s* = 2, 3, 4, respectively,

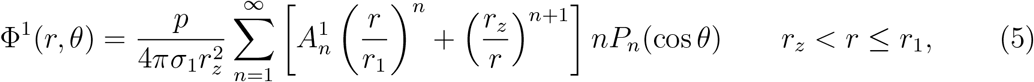

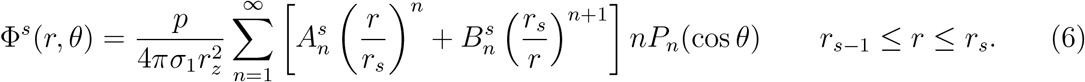

Here, Φ^*s*^(*r*), is the extracellular potential measured at location *r* in shell number *s*, of external radius *r_s_*, from current dipole moment *p* located at *r_z_*. The conductivity of sphere *s* is denoted by *σ_s_*, 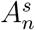 and 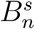 are constants depending on the shell radii and conductivities, and *P_n_*(cos*θ*) is the *n*-th Legendre Polynomial where *θ* is the angle between *r* and *r_z_*. From the boundary conditions listed in Equations (2)–(4), we can compute 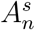, for *s* = 1, 2, 3, 4 and 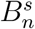, for *s* = 2, 3, 4, using the notation *σ_ij_* ≡ *σ_i_*/*σ_j_* and *r_ij_* ≡ *r_i_*/*r_j_*:

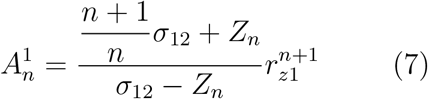

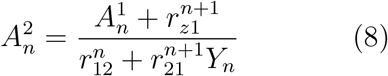

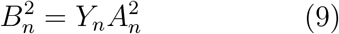

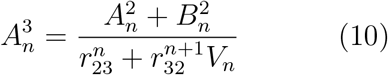

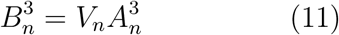

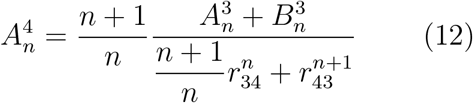

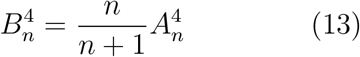

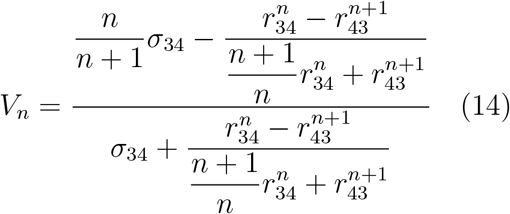

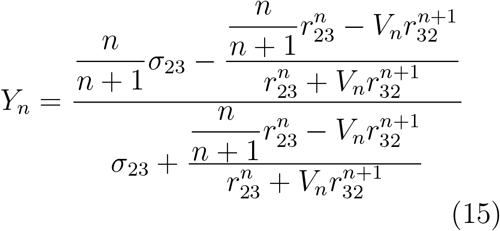

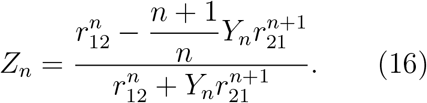

Equations (5) and (6) are in accordance with Equations (G.1.9–10) in Appendix G of Nunez and Srinivasan (2006) and Equation (A–1) in Srinivasan et al. (1998), Appendix A. However, some of the above constants (Equations (7)–(16)) are different from the ones given in Nunez and Srinivasan (2006) and Srinivasan et al. (1998), see Appendix A for specifics.

#### 2.2.2 Tangential dipole

The extracellular potential from a tangential dipole in a concentric-shells model is given by Equation (H.2.1) in Appendix H of Nunez and Srinivasan (2006), and takes the following from:

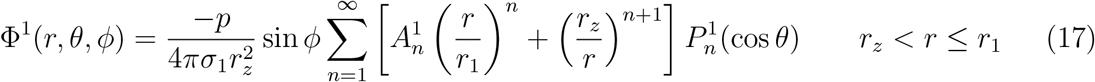

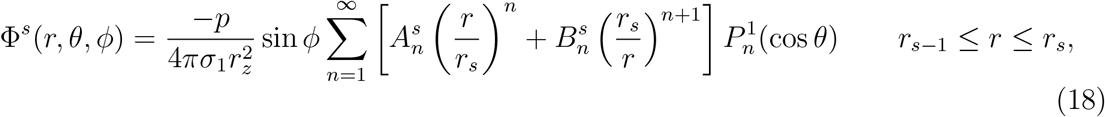

where *ϕ* is the azimuth angle and 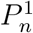 is the associated Legendre polynomial. When solving for the boundary conditions, Equation (2)-(4), we find that the constants 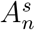 and 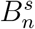 are the same as for the radial dipole solution, see Section 2.2.1.

In the results section we compare our analytical solution and the FEM simulations with the two published formulas for the potential in the four-sphere model given in Appendices G and H in Nunez and Srinivasan (2006), and in Appendix A in Srinivasan et al. (1998). For comparison we also present the approximate solution provided in Appendix G.4 in Nunez and Srinivasan (2006). Note that two corrections were done to the model presented in Srinivasan et al. (1998) before comparison. First of all, the multiplication factor *p*/*σ*_1_ was inserted in Equation (A-1), necessary to give potentials in units of volts. Secondly, a superscript in Equation (A-8) was changed, such that the right-hand-side included 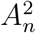 instead of 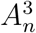, since this was obviously a typographical error. For more details on the different descriptions of the analytical four-sphere model, see Appendix A.

### 2.3 Finite Element Method

To find the numerical solution of the four-sphere model we solved the Poisson equation (Equation (1)) using the Finite Element Method (FEM). The first step was to construct a 3D numerical mesh representing the four-sphere head model geometry. We used the open-source program `gmsh` (Geuzaine, 2009), optimized using the `netgen` algorithm (Schöberl, 1997). Figure 1B shows the resulting mesh corresponding to the set of radii listed in Table 1. Note that our 3D FEM model-geometry implementation consists of five spheres: scalp, skull, cerebrospinal fluid (CSF), and two spheres together representing the brain tissue. However, the two innermost spheres (the innermost having a radius of 6 cm) are set to have the same conductivity, i.e., the value for brain tissue listed in Table 1. Thus, the model is effectively still a four-sphere model. We observed, however, that partitioning the four spheres into five and partitioning the inner sphere to a coarser mesh size reduced the overall mesh size and computational time while retaining the accuracy. The resulting mesh comprised of nearly 12.2 million tetrahedrons (2.1 million odd nodes) and we observed that at this resolution, the numerical results had converged.

The current sources were treated as point sources and the conductivity was set in each mesh point according to Table 1. Finally, the Poisson’s Equation and the boundary each mesh point according to Table 1. Finally, the Poisson’s Equation and the boundary conditions listed in Equations (2)–(4) were solved numerically with FEM. All FEM simulations were done with the open-source program `FEniCS` (Logg et al., 2012; Alnæs et al., 2015), with Lagrange P2 finite elements. The linear systems were solved by the the *Krylov Solver* employed with the *Conjugate Gradient* method, and the *Incomplete LU* factorization preconditioner.

### 2.4 Software

All the `Python` code for obtaining the potentials from a current dipole placed in a four-sphere head model using (i) the analytical formulation and (ii) the numerical method (FEM) are available under the GNU General Public License version 3 here: https://github.com/Neuroinflab/fourspheremodel. Additionally, the `Python` scripts to generate the figures presented in this manuscript are also included. We tested this code in Anaconda Scientific package on a Linux 64 machine. For easy uptake of this resource and verification, we provide the associated `conda` environment with all the specific libraries used to run this software, and a help file.

## 3 Results

### 3.1 Comparison between analytical and FEM results

EEG potentials were computed on the scalp surface with the analytical four-sphere model Φ(*r*_4_, *θ, ϕ*) and compared with the results from the FEM simulations for a current dipole **p**. To mimic a current dipole set up by cortical neurons, the dipole was placed in the brain layer (*s* = 1) of the four-sphere head model, close to the CSF boundary, cf. Figure 2A, E, I. We found that the analytical and FEM models gave similar results for both radial and tangential dipoles: the absolute value of the difference was more than two orders of magnitude smaller than the EEG signal itself for all dipole orientations (Figure 2).

**Figure 2:**
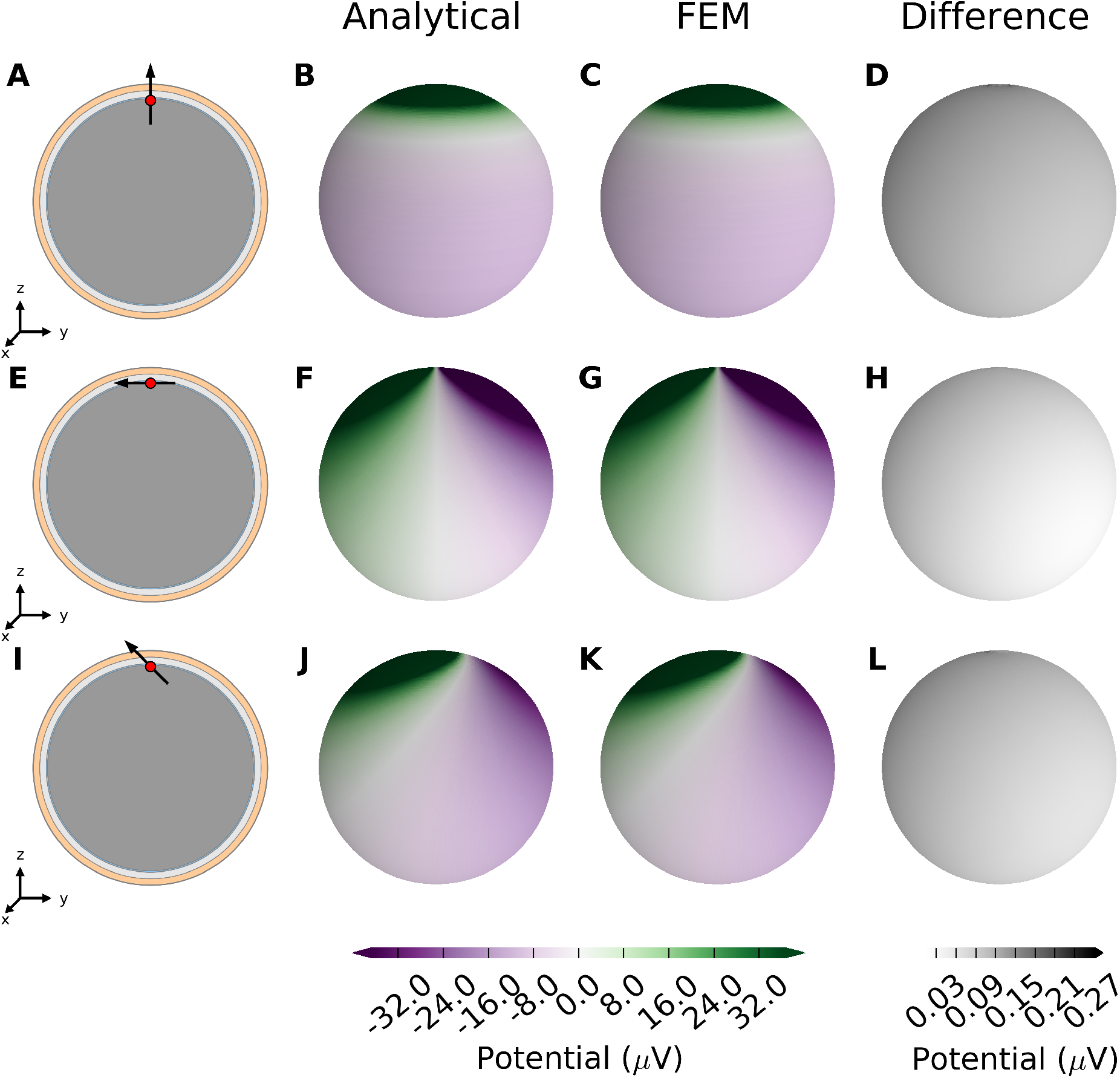
EEG potentials computed with four-sphere model and FEM simulation for radial, tangential, and 45-degree dipole. **(A)** A radial current dipole placed in the brain in the model as described in Table 1. The dipole (black arrow) is located at **r**_*z*_ = [0, 0, 7.8 cm] (red dot) and has a magnitude 10^−7^ Am to give scalp potentials some tens of microvolts in magnitude, typical for recorded EEG signals. **(B)** Resulting scalp potential calculated with the analytical four-sphere model. **(C)** Scalp potential computed with FEM. **(D)** Absolute difference between results from analytical calculation and FEM. The second row, panels **E-H** are equivalent to the top row, however for a tangential dipole parallel to *y* axis, in the *x* = 0 plane. The bottom row, panels **I-L** are equivalent to the top row, however for a dipole that subtends 45-degrees to the *z* axis in the *x* = 0 plane.

A more detailed comparison of EEG potentials predicted by the analytical model and the FEM model is shown in Figure 3. Here the computed EEG signal from a radial current dipole is shown for increasing polar angle *θ* between the current dipole position vector **r**_*z*_ and the measurement position vector **r**. The sphere radii and conductivity values are consistent with Nunez and Srinivasan (2006) (Table 1). The curve for the analytical results (blue line) overlaps the FEM results (red dots). This figure also demonstrates that previously published formulas give incorrect predictions.

**Figure 3:**
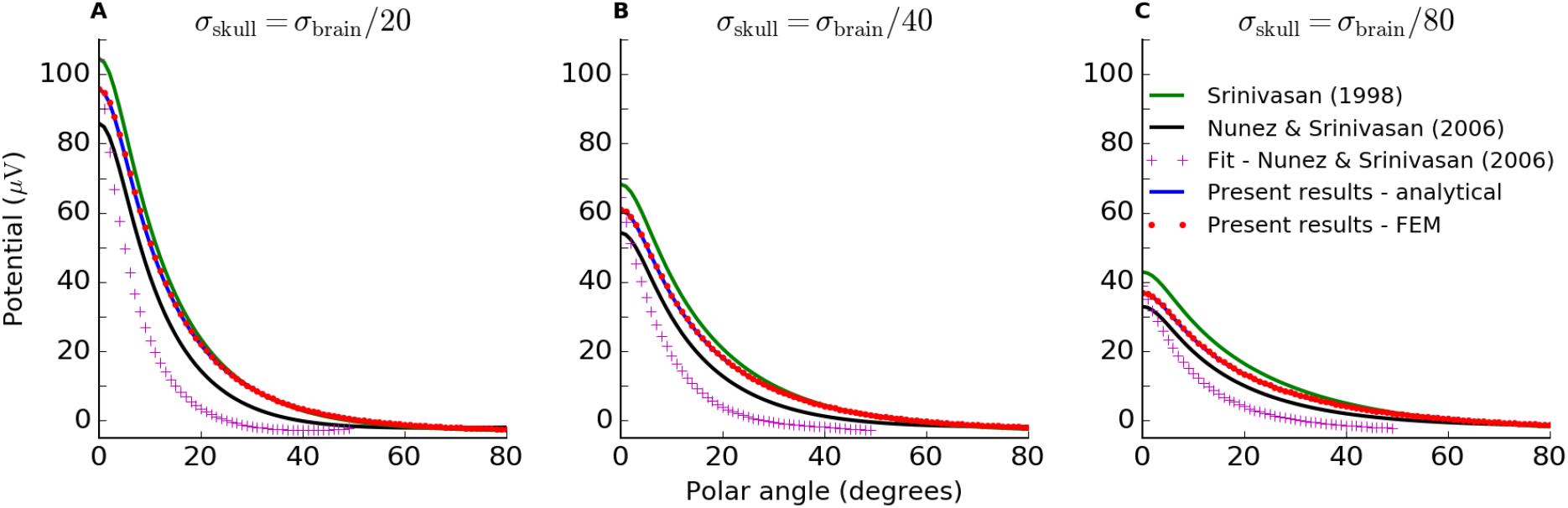
Analytical solution of four-sphere model matches FEM simulation. Scalp potentials from radial current dipole at position *r_z_* = 7.8 cm and magnitude 10^−7^ Am to give results in observable range, while still facilitiating direct comparison with the original plots in Srinivasan et al. (1998); Nunez and Srinivasan (2006). The resulting scalp potentials are shown for increasing polar angle *θ* between the current dipole and the measurement position vector. The different lines show calculations with the various formulations of the four-sphere model discussed in this paper, as well as the FEM simulation. The green line shows potentials obtained from Srinivasan et al. (1998), Appendix A, Equations (A-1 – 11). The black line shows results from applying the formulation given in Nunez and Srinivasan (2006), Appendix G, Equations (G.1.9–10) and (G.2.1–10). The approximate solution from Nunez and Srinivasan (2006), Appendix G.4, Equation (G.4.1–3) is given by the pink crosses. The analytical formulation of the four-sphere model presented here is shown in blue, and the FEM simulation is given by the red dots. Panels **A**, **B** and **C** show results for different values of the skull conductivity, i.e., *σ*_skull_ = *σ*_brain_/20, *σ*_brain_/40 and *σ*_brain_/80, respectively.

### 3.2 Limiting case

As an additional control we tested the limiting case where the conductivity was set to be the same for all four shells, i.e., *σ*_brain_ = *σ*_CSF_ = *σ*_skull_ = *σ*_scalp_, and equal to that of the brain (Table 1). In this case, the resulting scalp potentials should be the same as those calculated from a homogeneous single-sphere head model with radius equal to the scalp radius *r*_4_. For a dipole oriented along the radial direction inside a single homogeneous sphere, the surface potentials are given by Equation (6.7) in Nunez and Srinivasan (2006):

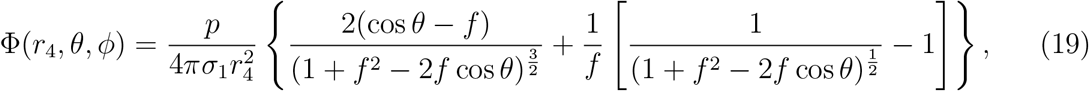

where *f* = *r_z_*/*r*_4_. Comparison between the simplified four-sphere models and the homogeneous single-sphere model showed perfect agreement for the present formulation, while the formulas listed in Srinivasan et al. (1998) and Nunez and Srinivasan (2006) give inaccurate predictions (Figure 4).

**Figure 4:**
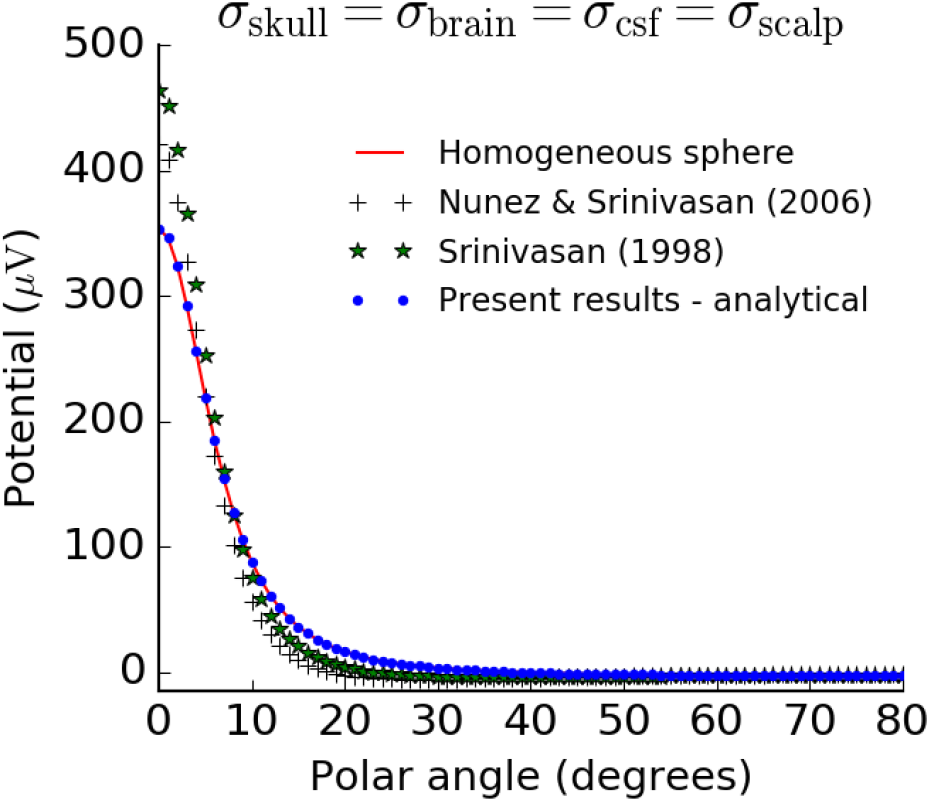
Analytical solution of the four-sphere model satisfies control test for limiting case. Four-sphere models in the limiting case where the conductivity of the skull, CSF, and scalp are equal to the conductivity of the brain, compared to the equivalent model for a single homogeneous sphere, Equation (19). We used a radial dipole of magnitude 10^−7^ Am positioned a distance *r_z_* = 7.8 cm away from the center of the sphere, consistent with Figure 2 and 3.

## 4 Discussion

In this note we have revisited the analytical four-sphere model for computing EEG potentials generated by current dipoles in the brain. The main contributions of this paper are the presentation of corrected and validated formulas, as well as numerical scripts for using them, allowing users to readily apply this important forward-model in the field of EEG analysis.

We also provide a set of FEM scripts which model the four-sphere model consistent with the analytical solution.

In addition to facilitating the use of the four-sphere model in EEG signal analysis (see, e.g., Peraza et al. (2012); Wong et al. (2008); Chu et al. (2012)), the present formulas and scripts will also be a resource for benchmarking comprehensive numerical schemes for computing EEG signals based on detailed head reconstructions such as the Finite Element Method (FEM) (Larson and Bengzon, 2013). The FEM approach is not restricted to specific head symmetry assumptions and can take into account an arbitrarily complex spatial distribution of electrical conductivity representing the electrical properties of the head. This is done by constructing a complicated numerical mesh for the head, a task that is often technically challenging. The present validated analytical solution for the four-sphere model can thus serve as a ground-truth benchmark against which the correctness and computational precision of such comprehensive numerical implementations can be tested.

## Acknowledgments, funding and conflict of interest statement

We thank Paul L. Nunez and Ramesh Srinivasan for useful personal contact. The study received funding from the Simula-UCSD-University of Oslo Research and PhD training (SUURPh) program, funded by the Norwegian Ministry of Education and Research, the European Union Horizon 2020 Research and Innovation Programme under Grant Agreement No. 720270 [Human Brain Project (HBP) SGA1], from the EC-FP7-PEOPLE sponsored NAMASEN Marie-Curie ITN grant 264872 and the Polish National Science Centre’s OPUS grant (2015/17/B/ST7/04123). The authors declare no conflict of interest.

## A Mathematical derivation of four-sphere model

The four-sphere model equations for radial and tangential dipoles are given in Equations (5), (6), (17) and (18). Here we describe how the seven unknown constants (Equation (7)–(16) can be determined by the seven boundary conditions (Equations (2)–(4)). We show the calculations for radial dipoles only, however, the derivation presented applies to both radial and tangential dipoles, due to similarity of the models.

We start by finding the derivative of Φ^s^(*r*; *θ*) from Equation (6):

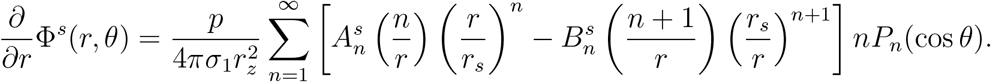

For the Neumann boundary condition on the scalp boundary, Equation (4), we make use of the relation above, and get:

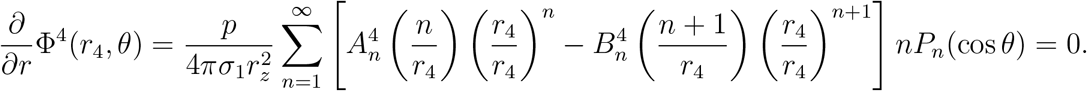

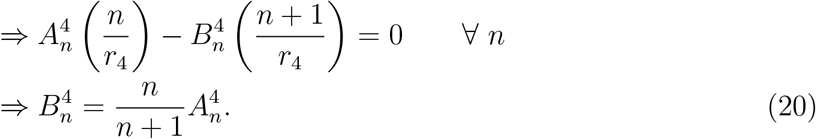

Next, we apply the Dirichlet boundary condition on the skull boundary, i.e., Equation (2) for *s* = 3:

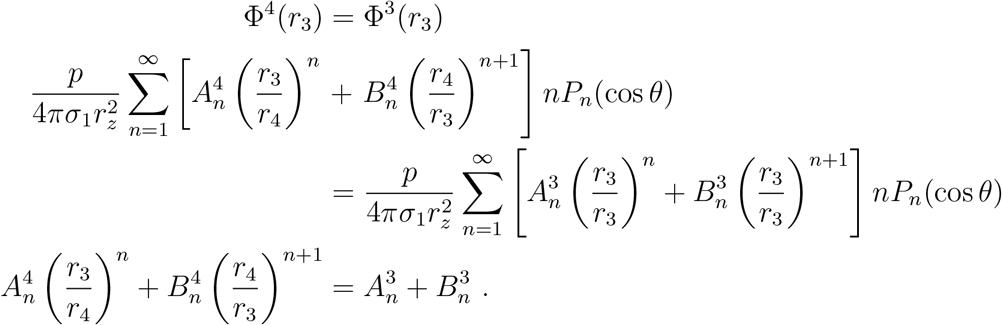

Inserting the expression for 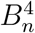, Equation (20), using the notation *r_i_*_*j*_ ≡ *r*_*i*_/*r_j_*:

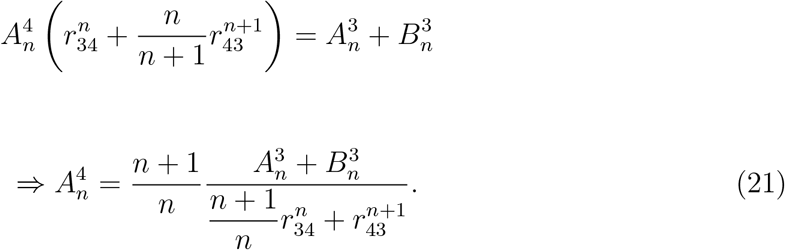

Note that the multiplication factor 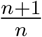 is missing in Nunez and Srinivasan (2006), Appendix G, Equation (G.2.9).

Further, we look at the Neumann boundary condition on the skull boundary, i.e. Equation (3) for *s* = 3, using the notation *σ_ij_* ≡ *σ_i_*/*σ_j_*:

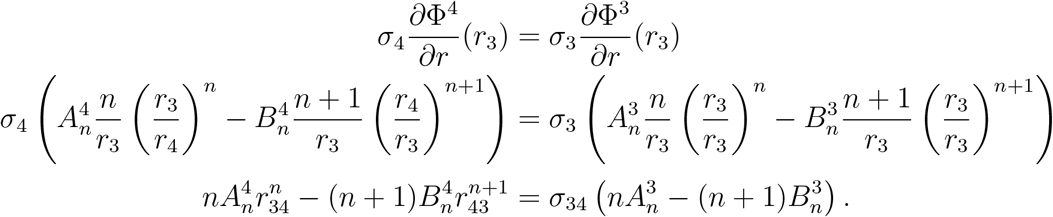

Inserting Equation (20),

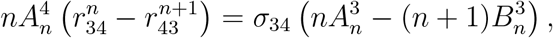

and applying Equation (21),

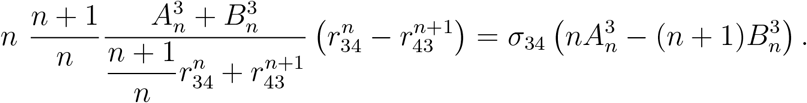

From this we find that,

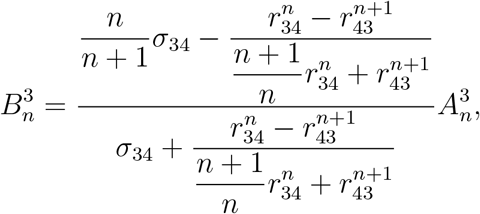

which we can write as:

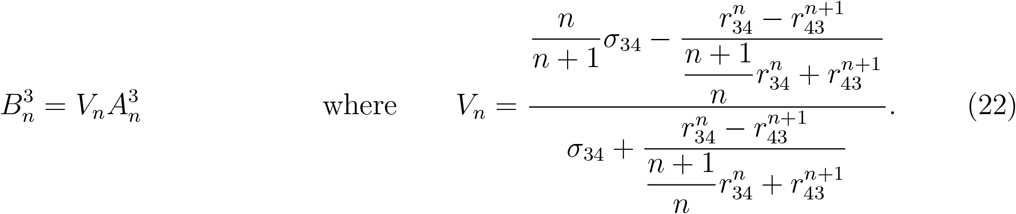

Here, the *σ*_34_-term in the numerator of *V_n_* differs from Nunez and Srinivasan (2006) (Equation (G.2.1)) and Srinivasan et al. (1998) (Equation (A-2)) in the sense that the multiplication factor is inverted.

For the CSF Dirichlet boundary condition we can follow the same procedure as for the skull Dirichlet boundary condition, and we get,

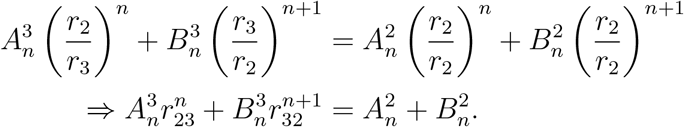

Inserting the expression for 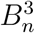 from Equation (22):

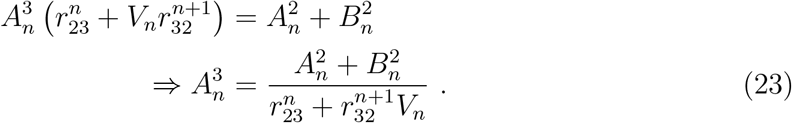

Here, we notice a typographical error in the expression for 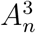 in Srinivasan et al. (1998), Equation (A-8): there should be an 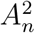-term in the numerator, not 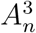.

Next, we apply the Neumann CSF boundary condition. Starting out with,

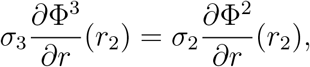

and making use of the expressions for 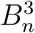 and 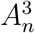, we find that,

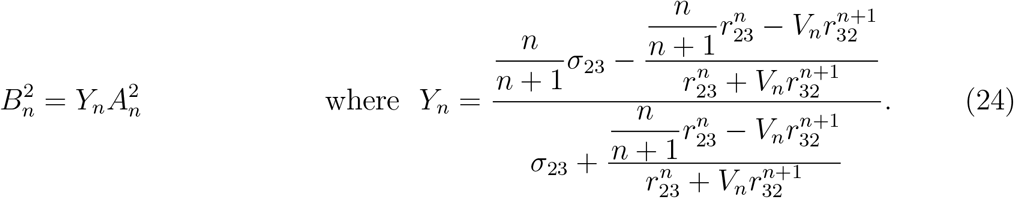

Note that there’s a subtle difference between the *Y_n_* presented here, and Nunez and Srinivasan (2006) (Equation (G.2.2)) and Srinivasan et al. (1998) (Equation (A-3)): The second term of the numerator is a fraction. Here, the 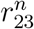 factor should not be multiplied by the whole fraction, but rather only the 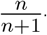-term in the numerator.

The Dirichlet boundary condition on the brain boundary is:

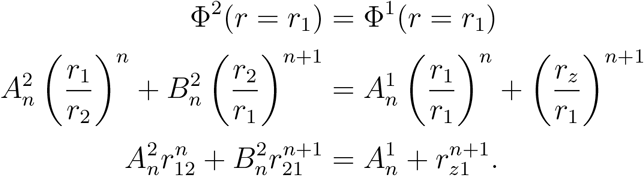

Inserting the expression for 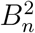 from Equation (24):

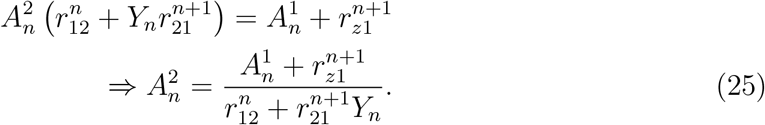

Finally, we solve the Neumann boundary condition on the brain boundary,

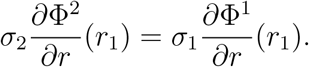

Inserting the expressions for 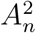 and 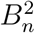 from Equations (25) and (24), we find,

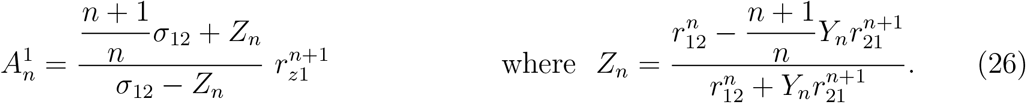

The 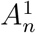-term in Srinivasan et al. (1998) (Equation (A-5)) is not consistent with Nunez and Srinivasan (2006) (Equation (G.2.4)) equal to Equation (26): a multiplication factor *p*/*σ*_1_ is lacking, 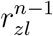 should be 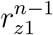. Moreover, 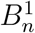 needs to be defined in order for the model description in Srinivasan et al. (1998), Appendix A to give potentials in brain tissue.

